# Structural characterization of the ACDC domain from ApiAP2 proteins of the malaria parasite

**DOI:** 10.1101/2024.02.09.579679

**Authors:** Marine Le Berre, Thibault Tubiana, Philippa Reuterswärd Waldner, Noureddine Lazar, Ines Li de la Sierra, Joana Mendonca Santos, Manuel Llinás, Sylvie Nessler

## Abstract

The Apicomplexan AP2 (ApiAP2) proteins are the best characterized family of DNA-binding proteins in the malaria parasite. Apart from the AP2 DNA-binding domain, there is little sequence similarity between ApiAP2 proteins and no other functional domains have been extensively characterized. One protein domain, which is present in a subset of the ApiAP2 proteins, is the conserved AP2-coincident domain mostly at the C-terminus (ACDC domain). Here we solved for the first time the crystal structure of the ACDC domain from two distinct *Plasmodium falciparum* ApiAP2 proteins and one orthologue from *P. vivax*, revealing a non-canonical four-helix bundle. Despite little sequence conservation between the ACDC domains from the two proteins, the structures are remarkably similar and do not resemble that of any other known protein domains. Due to their unique protein architecture and lack of homologues in the human genome, we performed *in silico* docking calculations against a library of known antimalarial compounds and we identified a small molecule that can potentially bind to any Apicomplexan ACDC domain within a pocket highly conserved amongst ApiAP2 proteins. Inhibitors based on this compound would disrupt the function of the ACDC domain and thus of the ApiAP2 proteins containing it, providing a new therapeutic window for targeting the malaria parasite and other Apicomplexans.

## Introduction

Malaria is an infectious disease caused by intracellular Apicomplexan parasites from the genus *Plasmodium*. The two most predominant *Plasmodium* species are *P. falciparum* and *P. vivax*, which annually kill over half a million people worldwide (WHO). The parasite lifecycle involves developmental stages in both the human and the *Anopheles* mosquito hosts, and mosquitos also serve as the vector for transmission. Malaria parasite development relies on the expression of specific sets of parasite genes at precise times throughout the lifecycle. Transcriptional regulation is largely coordinated by the Apicomplexa *Apetela*2 (ApiAP2) protein family, the largest and most well-characterized family of DNA-binding proteins in *Plasmodium* (Painter *et al*., 2011; Balaji, 2005; Jeninga *et al*., 2019). ApiAP2 proteins regulate transcription through sequence-specific interactions determined by small, 60 amino acid AP2 DNA-binding domains. These are found only in the protist ApiAP2 proteins and in the AP2/ERF family of plant transcription factors (Balaji, 2005; Feng *et al*., 2020). ApiAP2 proteins are encoded by all apicomplexan parasites and have been shown to play essential roles in transcriptional activation or repression throughout parasite development (Jeninga *et al*., 2019; Zarringhalam *et al*., 2023). Due to their essentiality and lack of mammalian orthologues, ApiAP2 proteins are considered as putative drug targets (Russell *et al*., 2022).

ApiAP2 proteins vary greatly in size (from 200 to 4109 amino acids in *P. falciparum*) and may contain 1 to 3 AP2 DNA-binding domains (Jeninga *et al*., 2019). Another domain of unknown function that is almost exclusively found in the ApiAP2 proteins is the AP2-Coincident Domain mostly at the C-terminus (ACDC) (Oehring *et al*., 2012). In *P. falciparum*, the ACDC domain is present only once in each protein and it is generally located at the proteins C-terminus, except for AP2-I, in which it is N-terminal (Oehring *et al*., 2012; Jeninga *et al*., 2019; Santos et al 2017). All Apicomplexa encode ApiAP2 proteins containing ACDC domains, although only a subset of ApiAP2 proteins encode the domain. In *P. falciparum*, for instance, 8 out of the 27 ApiAP2 proteins have an ACDC domain (Oehring *et al*., 2012). The ACDC domain is characterized by a ∼90 residues long consensus sequence that is poorly conserved between ApiAP2 proteins (**Sup. Figure 1**).

In this study, we solved and compared the crystal structures of two distinct ApiAP2 ACDC domains from *Plasmodium*: (i) the C-terminal ACDC of AP2-O5 and (ii) the N-terminal ACDC domain of AP2-I. AP2-O5 has been shown in *P. berghei* to regulate the expression of genes encoding for proteins implicated in the mobility of the ookinete, the mosquito midgut invasive stages of the parasite, and it has been reported to be an essential protein for asexual blood-stage development of *P. berghei* and *P. falciparum* but not *P. yoelli* (Bushell *et al*., 2017; Zhang *et al*., 2017; Zhang *et al*., 2018). The PfAP2-I transcription factor regulates expression of proteins allowing invasion of the host erythrocytes in *P. falciparum* (Santos *et al*., 2017) and it is essential for survival of all *Plasmodium* species (Jeninga *et al*., 2019).

Our structural analysis reveals that the ACDC domain adopts a novel conserved fold characterized by a left-handed orthogonal 4-helix bundle. We also performed *in silico* docking calculations, which suggested that compounds from the Tres Cantos Anti-Malarial Set (TCAMS) (Gamo *et al*., 2010) bind to the ACDC domain. Taken together, these results demonstrate the druggable potential of the ACDC domains not only from *Plasmodium* but also from other apicomplexan parasites containing ApiAP2 proteins with ACDC domains, like *Toxoplasma gondii*, responsible for toxoplasmosis, or *Cryptosporidium*, the main cause of parasite-derived diarrhea in the world.

## Materials & Methods

### Cloning

Synthetic genes of full-length AP2-I from *P. falciparum* (UniProt entry Q8IJW6; ORF name PF3D7_1007700; 1597 amino acids) and *P. vivax* (UniProt entry A0A564ZT73; ORF name PVP01_0807400; 1397 amino acids) were codon optimized for expression in *Escherichia coli* (ThermoFisher). The gene coding for the AP2-O5 protein of *P. falciparum* (UniProt entry Q8IKY0; ORF name PF3D7_1449500; 715 amino acids) was amplified from genomic DNA prepared from *P. falciparum* parasites using the DNeasy Kit (Qiagen) according to the manufacturer’s instructions. Gene fragments coding for residues 1-152 of PfAP2-I and residues 30-230 of PvAP2-I were amplified by PCR using the primers 3 and 4 or 5 and 6 (Eurofins), respectively (**Sup. Table 1)**, and then cloned into a pET28 plasmid (Novagen) with T4 DNA ligase (Thermo Scientific) using the restriction enzyme pairs NcoI/XhoI (Thermo Scientific) to generate C-terminal Strep Tag II affinity tags. The gene fragment coding for residues 625-715 of PfAP2-O5 was amplified by PCR using the primers 1 and 2 (**Sup. Table 1**), and then cloned into a pSL1045 plasmid (Addgene) using XhoI and NcoI (New England Biolabs) followed by T4 DNA ligase (New England Biolabs) to generate a C-terminal His-tag preceded by a TEV protease cleavage site. The three plasmids were transformed into ultra-competent *E. coli* DH5α bacteria (ThermoFisher). Correct assembly was verified by PCR using Taq 2X Master Mix (New England BioLabs) and Sanger sequencing (Genewiz).

### Proteins expression and purification

Expression of the ACDC domains of PfAP2-O5, PfAP2-I and PvAP2-I was carried out in *E. coli* (DE3)-Gold strains in 800 ml of 2YT medium at 37°C for 3 hours. Induction was done by adding 0.5 mM IPTG (Sigma). Cells were harvested by centrifugation (4,500 G for 20 min at 8°C), resuspended in 40 ml of buffer A (20 mM Tris-HCl pH 7.5, 200 mM NaCl) and stored at −20°C. After thawing, a protease inhibitor cocktail (Thermo Scientific) was added to the resuspended bacterial pellet and cell lysis was carried out by sonication (probe-tip sonicator Branson). After centrifugation (20,000 G for 30 min at 8°C), the 40 ml soluble fractions were purified by affinity chromatography. Strep-tagged AP2-I ACDC domains were loaded onto a 5 ml StrepTrap HP column (Cytiva) equilibrated at 4°C in buffer A for FPLC chromatography. After extensive washing with buffer A, the Strep-tagged proteins were eluted with buffer A supplemented with 2.5 mM desthiobiotin. The His-tagged AP2-O5 ACDC domain was loaded onto a 3 ml NTA-Ni resin on a benchtop column, washed extensively with cold buffer A and then with cold buffer A supplemented with 20 mM imidazole. Finally, the AP2-O5 ACDC domain was eluted with cold buffer A containing 400 mM imidazole. The purified fractions were then loaded onto a Hiload 16/60 Superdex 75 prep grade size exclusion chromatography column equilibrated in buffer A to eliminate aggregates. Protein concentrations were determined by measurement of the absorbance at 280nm using a NanoDrop spectrophotometer (Thermo Fisher). Purified proteins were concentrated using Vivaspin 5,000 MWCO concentrators, aliquoted and stored at −80°C in buffer A.

### Crystal structure determination and analysis

Crystallization assays were performed at 18°C in 96-well plates using commercial crystallization screening kits (Molecular Dimension) in 200 nl droplets (100 nl protein + 100 nl reservoir solution) prepared by using the Cartesian Microsys pipetting robot (PROTEIGEN) at the 12BC crystallization facility. Prior to diffraction assays, crystals were cryoprotected by transferring them into droplets containing their crystallization solution supplemented with 30% PEG 400 and flash-frozen in liquid nitrogen. Diffraction data were collected at the synchrotron SOLEIL (Saint-Aubin, France) on beamline Proxima-2. Data processing was performed using the XDS package (Kabsch, 2010). Phasing was performed by molecular replacement using the PHASER program (McCoy *et al*., 2007). Initial models were predicted using ColabFold v1.5.3: AlphaFold2 using MMseqs2 (Jumper *et al*., 2021; Mirdita *et al*., 2022). For the ACDC domain of PfAP2-O5, AlphaFold2 predicted 5 similar models (rmsd below 0.5Å) with high confidence scores (predicted Local Distance Difference Test (pLDDT) > 90) and the rank-1 model was efficiently used for phasing by molecular replacement. For the ACDC domain of PfAP2-I and PvAP2-I, disordered regions of the AlphaFold2 models were deleted for phasing.

Protein structure refinement was performed using PHENIX (Liebschner *et al*., 2019) and further improved by iterative cycles of manual rebuilding using COOT (Emsley *et al*, 2010). The final structures were deposited into the Protein Data Bank (PDB) (Berman *et al*, 2000). We used PyMOL (The PyMOL Molecular Graphics System, version 1.2r3pre, Schrödinger, LLC n.d.) for graphic analysis of the structures and preparation of figures. With used ESPRIPT (Gouet *et al*., 1999) for graphical enhancement of sequence alignments. Protein assemblies were analyzed using the PDBePISA server at the European Bioinformatics Institute (EBI) (http://www.ebi.ac.uk/pdbe/prot_int/pistart.html) (Krissinel & Henrick, 2007). Protein structure comparison was performed using FoldSeek with the 3Di/AA mode (https://search.foldseek.com) (Van Kempen *et al*., 2023) against the ∼200 thousands experimental structures of the Protein Data Bank (Berman *et al*, 2000) and the ∼200 million predicted structures of the AlphaFold2 database (Varadi *et al*., 2022).

### *In silico* docking screen

The 3 crystal structures solved in this study were prepared for docking simulations with the MGLTool’s (Morris *et al*., 2009) using prepare_receptor.py script. A screening box surrounding a conserved region was used as target for the docking calculation. We screened against the 13,533 small molecules from the Tres Cantos Anti-Malarial Set (TCAMS), which have all been shown to inhibit *P. falciparum* growth by at least 80% at 2 μM concentration (Gamo *et al*., 2010). The platform KNIME (P. Mazanetz *et al*., 2012) was employed to process the SMILEs notation of each compound. The molecules were desalted using the vernalis node and protonated prior to transformation into their 3D forms. The latter were minimized using the MMFF94 force field over 2000 iterations, and Gasteiger partial charges (Gasteiger and Marsili, 1980) were assigned via RDKIT (https://www.rdkit.org/, n.d.) nodes. The refined molecular library was then stored in a single SDF file. Torsional angles for each molecule were calculated using the Meeko preparation script (https://github.com/forlilab/Meeko), resulting in the assembly of individual ligands into distinct PDBQT files. During this process, macrocyclic components were treated as conformationally rigid. The docking protocol was executed using SMINA (Koes *et al*., 2013) with the Vinardo scoring function (Quiroga and Villarreal, 2016) set to explore 20 binding modes and an exhaustiveness parameter of 8 with 8 cores per molecule. Screening results were parsed and analyzed using in-house Python notebooks.

### Molecular Dynamic Simulation protocol

Prioritized compounds were prepared for molecular dynamics (MD) simulations. The parameterization of these compounds was conducted utilizing ACPYPE (Sousa Da Silva and Vranken, 2012), which serves as a bridge to Antechamber (Wang *et al*., 2004). The selection of initial conformations was guided by visual assessments aimed at optimizing interactions with the targeted pocket. Molecular dynamics simulations were performed with Gromacs 2022 (Abraham *et al*., 2015) and the Amber 99SB-ILDN force field (Lindorff-Larsen *et al*., 2010) on GPU. The complexes were solvated with TIP3P water model (Jorgensen *et al*., 1983) within a cubic periodic box, with a minimum distance of 1.5 nm from the protein-ligand complex to the edge of the box. Charge neutrality was achieved by the addition of appropriate Na^+^ or Cl^-^ ions. Initial energy minimization was carried out over 1000 steps using a conjugate gradient algorithm, integrating a steepest descent every 50 steps. This was followed by a heating phase over 500 ps, where the temperature was increased from 0 to 300K under NVT conditions (constant temperature, constant volume). The Berendsen thermostat (Berendsen *et al*., 1984) was applied with a time constant of 0.1 ps, while positional restraints were maintained on protein and heavy atoms of the ligand, exerting a force constant of 1000 kJ.mol.nm^2^. Subsequently, the systems underwent pressure equilibration for 500 ps in NPT conditions (constant temperature, constant pressure) employing the Berendsen barostat, set with a time constant of 1 ps and a compressibility factor of 4.5×10^−5^ bar^-1^, maintaining the aforementioned positional restraints. Restraints were removed for production and each system was simulated over 100 ns. The integration timestep was fixed at 2 fs for both equilibration and production phases. The LINCS algorithm (Hess, 2008) was employed to constrain bonds between heavy atoms and hydrogens. The Verlet cut-off scheme (Páll and Hess, 2013) was adopted for managing long-range interactions, with a threshold distance of 1 nm. We used the Protein-Ligand Interaction Profiler PLIP (Adasme *et al*., 2021) for the analysis of non-covalent interactions between the ACDC domains and the selected compounds.

## Results

### Crystal structure of the C-terminal ACDC domain of AP2-O5

The ACDC domain of PfAP2-O5 (residues 625-715) was efficiently expressed and purified from *E. coli*. The best crystals were obtained in 30% PEG400, 0.1 M MES pH 6.5, 0.1 M sodium acetate and diffracted up to 1.75Å resolution in space group P2_1_2_1_2 with two molecules per asymmetric unit (**Table 1**). We found that the PfAP2-O5 ACDC domain forms a left-handed orthogonal 4-helix bundle with a break in the middle of helix α3 (**Figure 1A**). The refinement procedure yielded a structure characterized by a R_work_ of 20.98% and a R_free_ of 24.78% (**Table 1**). The two polypeptide chains of the asymmetric unit display the same fold (rmsd of 0.61Å over 87 aligned Cα atoms). Chain B is fully visible (residues 625-715). In chain A, 5 amino acids (ENLYF) from the TEV site introduced between the protein sequence and the C-terminal His-tag form an additional turn in the C-terminal helix α4 whereas residues 676-677 from loop α2-α3 are not visible in the electron density map. Analysis of the protein interfaces in the crystal packing revealed that two molecules of the asymmetric unit display a buried area of 2570Å^2^, a solvation free energy gain ΔG^int^ of −22.1 kcal/mol and a Complex formation Significance Score of 1, suggesting that they may form a stable dimer in solution (**Figure 1B**). However, the isolated domain eluted as a monomer from the SEC purification column (**Sup. Figure 2**). In addition, given that the first 625 residues of PfAP2-O5 are missing in the structure of the ACDC domain, it is unclear if the ACDC domain dimerizes in the context of the full-length protein.

**Table 1:** X-ray diffraction data processing and refinement statistics.

|  | **PfAP2-I**<br>(PDB ID 8RXO) | **PvAP2-I**<br>(PDB ID 8RWU) | **PfAP2-O5**<br>(PDB ID 8RXA) |
| --- | --- | --- | --- |
| Crystallization condition | 1.5 M Ammonium sulfate,<br>0.1 M HEPES pH 7.5, 2%<br>Peg400 | 30% Peg400, 0.1 M MES<br>pH 6.5, 0.1 M Sodium<br>acetate | 30% Peg400, 0.1 M MES<br>pH 6.5, 0.1 M Sodium<br>acetate |
| **Data processing statistics** |  |  |  |
| Space group | P41 21 2 | P41 21 2 | P21 21 2 |
| Unit-cell parameters (Å) | 78.08; 78.08; 127.84 | 78.49; 78.49; 123.88 | 58.93; 108.93; 34.01 |
| Unit-cell parameters (°) | 90; 90; 90 | 90; 90; 90 | 90; 90; 90 |
| Resolution range (Å)* | 50.0 – 3.000 (3.18-3.00) | 50.0 – 3.150 (3.34-3.15) | 50.0 - 1.758 (1.86-1.75) |
| No. of unique reflections | 8421 (1303) | 7133 (1106) | 22718 (3510) |
| Completeness (%) | 99.9 (99.6) | 99.7 (98.7) | 99.5 (96.8) |
| CC<sub>1/2</sub> | 0.999 (0.71) | 1.0 (0.495) | 0.999 (0.941) |
| Mean I/σ(I) | 12.81 (0.86) | 21.53 (1.18) | 22.24 (2.23) |
| R<sub>meas</sub> (%)<sup>a</sup> | 19.8 (259.2) | 11.8 (308.4) | 5.9 (97.4) |
| **Refinement statistics** |  |  |  |
| Resolution range (Å) | 49.46 - 3.00 (3.23 - 3.00) | 48.62 - 3.15 (3.61 - 3.15) | 40.0 - 1.75 (1.79 - 1.75) |
| No. of molecules/a.u. | 2 | 2 | 2 |
| R<sub>work</sub> (%)<sup>b</sup> | 25.28 (39.67) | 24.07 (30.47) | 20.99 (32.62) |
| R<sub>free</sub> (%)<sup>c</sup> | 27.46 (50.84) | 27.81 (36.07) | 24.86 (36.75) |
| Ramachandran |  |  |  |
| Favored (%) | 91.40 | 98.24 | 98.90 |
| Outliers (%) | 0.00 | 0.59 | 0.00 |
| R.M.S.D. |  |  |  |
| Bond lengths (Å) | 0.004 | 0.001 | 0.014 |
| Bond angles (°) | 0.610 | 0.359 | 1.304 |
| Chirality | 0.040 | 0.035 | 0.065 |
| Planarity | 0.004 | 0.004 | 0.008 |
| Dihedral | 3.050 | 1.532 | 19.405 |
| Average B, all atoms (Å<sup>2</sup>) | 112.25 | 124.79 | 41.05 |

\* Numbers in parentheses represent values in the highest resolution shell.
<sup>a</sup> $R_{\text{meas}} = \sum_{\text{hkl}} [N/N-1]^{1/2} \sum_i |I_i(\text{hkl}) - \langle I(\text{hkl}) \rangle| / \sum_{\text{hkl}} \sum_i I_i(\text{hkl})$ where N is the multiplicity of a given reflection, $I_i(\text{hkl})$ is the integrated intensity of a given reflection and $\langle I(\text{hkl}) \rangle$ is the mean intensity of multiple corresponding symmetry-related reflections.
<sup>b</sup> $R_{\text{work}} = \sum ||\text{Fobs}| - |\text{Fcalc}|| / \sum |\text{Fobs}|$ , where $|\text{Fobs}|$ and $|\text{Fcalc}|$ are the observed and calculated structure factor amplitudes respectively.
<sup>c</sup> $R_{\text{free}}$ is the same as $R_{\text{work}}$ but calculated with a 20% subset of all reflections that was never used in refinement.

**Figure 1:**
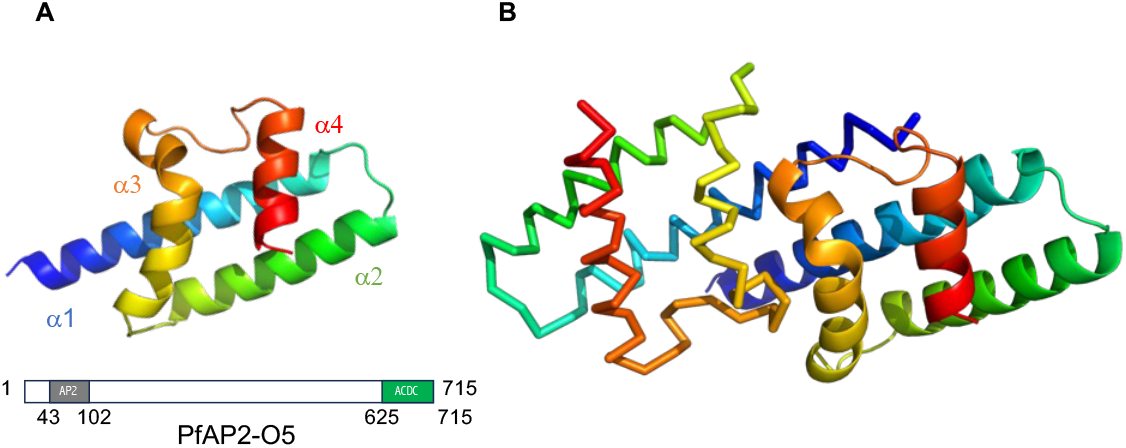
Crystal structure of the C-terminal ACDC domain of PfAP2-O5. **A** - The polypeptide chain is shown as cartoon colored by spectrum from blue (N-terminus) to red (C-terminus). **B** - The dimer observed in the symmetric unit is shown with one subunit represented as cartoon and the other as ribbon.

### Crystal structure of the N-terminal ACDC domain of AP2-I

The N-terminal ACDC domain of PfAP2-I (residues 60-150) and PvAP2-I (residues 59-149) are 92% identical. Several constructs containing the ACDC domain were tested for heterologous expression in *E. coli* and finally it was the extended fragments covering residues 1-152 from PfAP2-I and residues 30-230 from PvAP2-I that were efficiently expressed and purified from *E. coli* and used for crystallization assays.

The best crystals of the PfAP2-I ACDC domain were obtained in 1.5 M ammonium sulfate, 0.1 sodium HEPES pH 7.5, 2% PEG 400 and those of the PvAP2-I ACDC domain in 30% PEG 400, 0.1 M MES pH 6.5, 0.1 M sodium acetate. Both diffracted up to almost 3Å resolution in space group P4_1_2_1_2 with the same cell parameters (**Table 1**). As with the PfAP2-O5 ACDC domain, they each contained two molecules per asymmetric unit. AlphaFold2 proposed confident models for the core ACDC domain of PfAP2-I and PvAP2-I (residues 60-150) but the N- and C-terminal extensions of both elongated fragments were predicted as fully disordered. The proposed models were similar to those of the PfAP2-O5 ACDC domain but loop α3-α4 (residues 120-130), which was predicted with low accuracy according to the confidence pLDDT score, needed to be deleted for efficient phasing by molecular replacement. The final structure of PfAP2-I was refined at 3.0Å resolution with a R_work_ of 24.65% and a R_free_ of 26.35%, and that of PvAP2-I was refined to 3.15Å resolution with a R_work_ of 24.07% and a R_free_ of 27.81% (**Table 1**). With both fragments, electron density could only be observed for the core ACDC domain, resulting in an apparent solvent content of 73% in the crystal, and confirming that the N- and C-terminal extensions are disordered. Both AP2-I structures were highly similar with a root-mean-square deviation (RMSD) of 0.76Å over 171 superimposed Cα atoms of the 2 chains contained in the asymmetric unit (**Figure 2A**). Because they are indistinguishable, we will consider them as one, called Pf/PvAP2-I ACDC domain, in the rest of the manuscript. Electron density was clearly visible to reconstruct the loop α3-α4 omitted in the initial model (**Sup. Figure 3**) but it did not correspond to the conformation observed in the AlphaFold2 model and in the crystal structure of PfAP2-O5 (**Figure 1**). Instead, the AP2-I ACDC structure revealed a swapping of the C-terminal helix α4 between the two subunits of the asymmetric unit (**Figure 2A)**, resulting in a dimer characterized by a ΔG^int^ of −26.0 kcal/mol and a buried surface area of 2660 Å^2^. Interestingly, the PDBePISA analysis revealed two other stable interfaces (ΔG^int^ of −19.3 and −11.7 kcal/mol, respectively) in the AP2-I crystal packing but none of these three interaction modes corresponded to the dimer observed in the crystal structure of the PfAP2-O5 ACDC domain.

**Figure 2:**
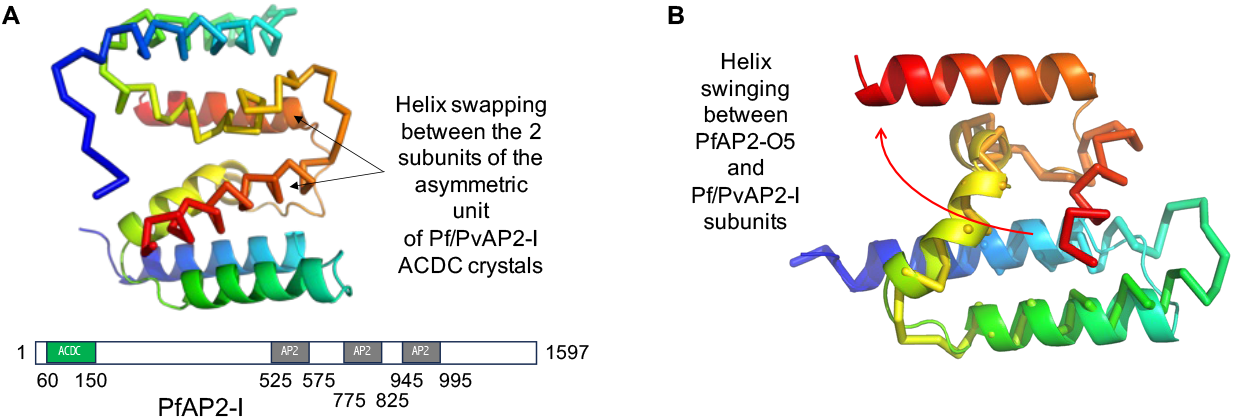
Crystal structure of the N-terminal ACDC domain of Pf/PvAP2-I. **A** - The dimer observed in the asymmetric unit is shown with one subunit represented as cartoon and the second as ribbon, both colored by spectrum. Swapping of helix α4 (in red) between the 2 subunits of the asymmetric unit is highlighted. **B** - Superimposition of the cartoon trace of one subunit of Pf/PvAP2-I with one subunit of PfAP2-O5 shown as ribbon. Both molecules are colored by spectrum. The swinging of helix α4 between the AP2-O5 and AP2-I conformations, is highlighted by an arrow.

### Structure analysis of the ACDC domain

Superimposition of the ACDC domains of Pf/PvAP2-I and PfAP2-O5 (**Figure 2B**) shows that, despite poorly conserved sequences (17.58% identity) (**Figure 3**), the ACDC domains of PfAP2-O5 and Pf/PvAP2-I display similar structures, except for the position of helix α4, which undergoes a large swinging motion between the two conformations. However, it is not clear if the C-terminal helix swapping observed in the dimeric crystal structure of the isolated N-terminal ACDC domain of AP2-I (**Figure 2A**) is present in the context of the full-length protein when helix α4 is followed by more than one thousand amino acids. In the absence of additional data to answer this question and the unknown biological relevance of the observed dimers, we focused our structural analysis on the PfAP2-O5 ACDC subunit, which matches the structure predicted by AlphaFold2.

**Figure 3:**
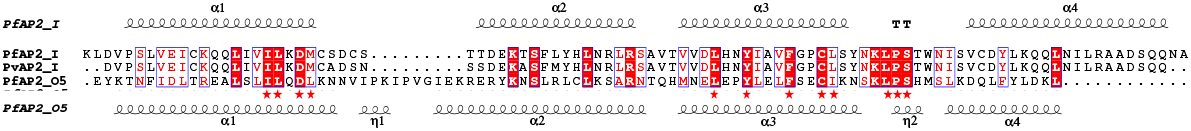
Multiple sequence alignment of the 3 ACDC domains characterized in this study. Secondary structure elements of PfAP2-I and PfAP2-O5 are shown at the top and the bottom of the alignment, respectively. Strictly and highly conserved residues are highlighted in white in red boxes and in red in white boxes, respectively. Residues involved in the characteristic α1/α3 interaction are highlighted by red stars.

A search for structural homologues of the PfAP2-O5 ACDC domain using Foldseek (Van Kempen *et al*., 2023) did not identified close experimental structures in the Protein Data Bank. Similar domains were only found among predicted models of proteins from various Apicomplexan species. The idiosyncratic kink observed in helix α3 of our three structures of ACDC domains is predicted to be present in all homologous ACDC domains found in ApiAP2 proteins, regardless of the Apicomplexan species (**Figure 4**) and can thus be considered the signature fold of the ACDC domain. These findings support the uniqueness of the ACDC structural domain characterized in this study.

**Figure 4:**
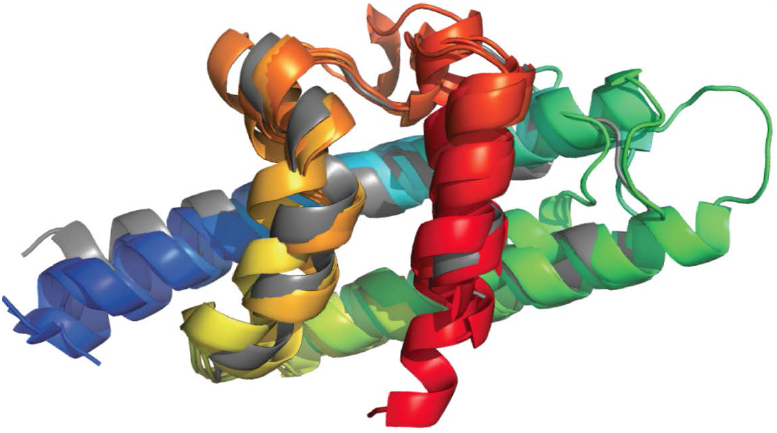
Superimposition of our crystal structure of the PfAP2-O5 ACDC and five randomly selected AlphaFold2 models of ACDC domains of ApiAP2 proteins from distinct Apicomplexa. PfAP2-O5 is shown in gray and the homologues are colored by spectrum (*Babesia ovata* A0A2H6K6P4, *Besnoitia besnoiti* A0A2A9MKU0, *Toxoplasma gondii* A0A7J6K727, *Theileria parva* Q4N6P4 and *Eimeria tenella* U6KJL8) (https://www.ebi.ac.uk/interpro/entry/pfam/PF14733/alphafold/#table).

### Docking analysis of plasmodial inhibitors

The kinked helix α3 of the ACDC domain is wrapped around helix α1, and this α1/α3 interaction involves residues conserved not only between PfAP2-I, PvAP2-I and PfAP2-O5 (**Figure 3**) but also across 45 aligned sequences of ACDC domains from distant apicomplexan homologues (https://www.ncbi.nlm.nih.gov/Structure/cdd/cddsrv.cgi?uid=434166). These conserved residues form a hydrophobic patch on the surface of the ACDC domain on the opposite side of the α4 swinging helix (**Figure 5A**). To explore ligands that could bind to any ACDC domain as potential inhibitors of ApiAP2 protein function by disrupting this domain, we used this conserved region to specify a screening box for *in silico* molecular docking (**Figure 5B** and **Sup. Table 2**). We screened the 13,533 small molecules of the Tres Cantos Antimalarial Set (TCAMS) (Gamo *et al*., 2010) against the 3 ACDC domain structures solved in this study. All protein side chains were fixed in a rigid conformation, except for 3 residues of the targeted pocket (Q71, L119, and N122 in PfAP2-I), which were designated as flexible to accommodate ligand binding. The best hits were selected based on the lowest SMINA docking energy scores, lowest RMSD from the top-ranked poses (**Figure 6A**) and proximity to the targeted conserved pocket. This proximity was quantified by the shortest distance from any ligand atom to the geometric center of specific residues (Q71, C118, N122 in PfAP2-I; C68, Q71, L119 in PvAP2-I; L712, L715, I694 in PfAP2-O5). The typical affinity score hovered around −6.7 kcal/mol (**Sup. Figure 4)**. Our search prioritized compounds demonstrating consistently low energy scores across the 3 ACDC domains. Out of the over 13,000 compounds, only seven met our stringent selection criteria – TCMDC-124223, TCMDC-124284, TCMDC-125666, TCMDC-134543, TCMDC-137354, TCMDC-139888 and TCMDC-140881 (**Table 2**). The compounds TCMDC-124223 and TCMDC-125666 are highly similar, with a single missing bond. TCMDC-124284 and TCMDC-137354 are also chemically related with a methyl group replaced by an oxygen atom.

**Table 2:**
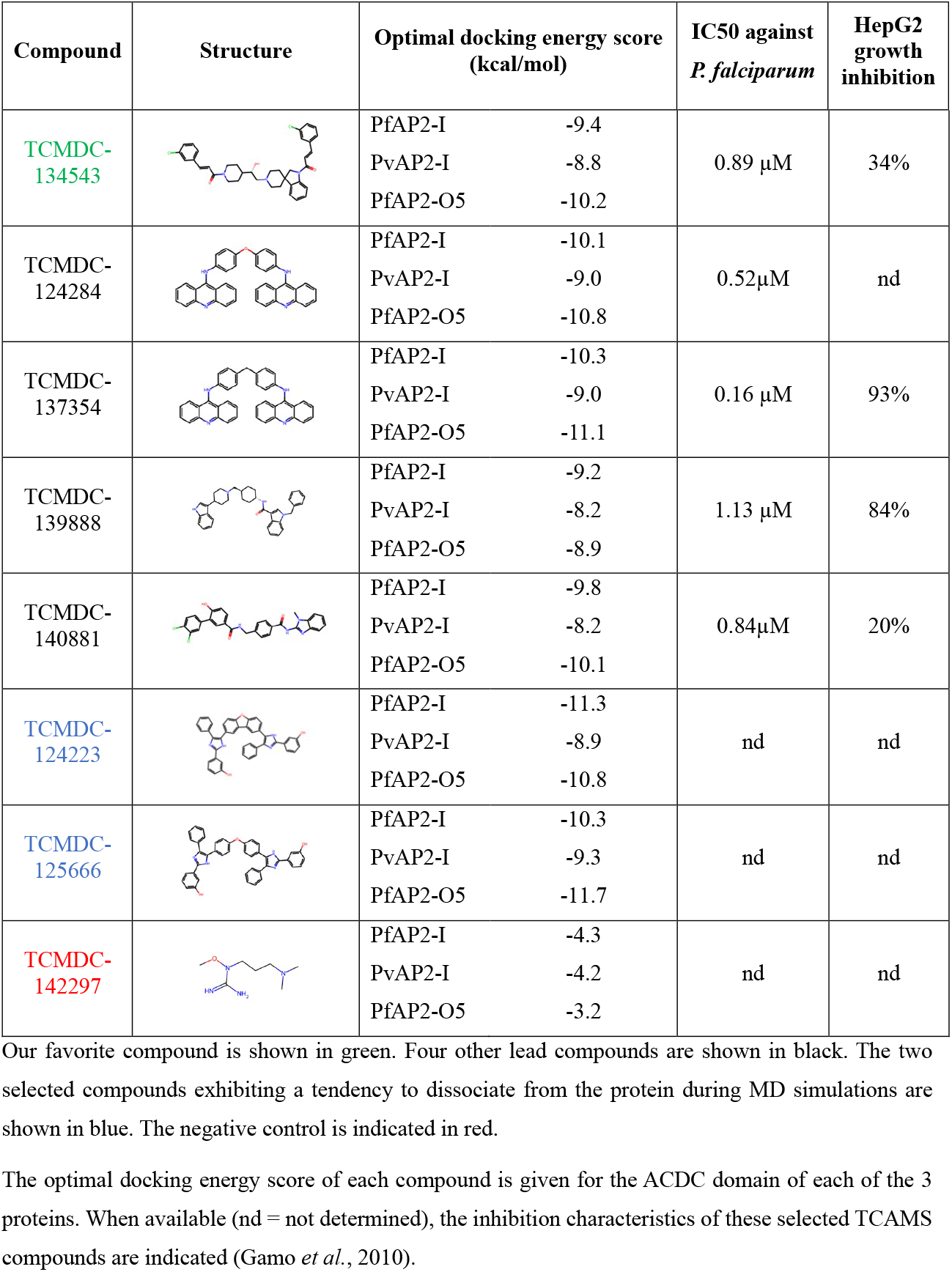
Structure of the selected compounds of the TCAMS.

| Compound | Structure | Optimal docking energy score (kcal/mol) | IC50 against *P. falciparum* | HepG2 growth inhibition |
| --- | --- | --- | --- | --- |
| TCMDC-134543 | | PfAP2-I -9.4<br>PvAP2-I -8.8<br>PfAP2-O5 -10.2 | 0.89 μM | 34% |
| TCMDC-124284 | | PfAP2-I -10.1<br>PvAP2-I -9.0<br>PfAP2-O5 -10.8 | 0.52 μM | nd |
| TCMDC-137354 | | PfAP2-I -10.3<br>PvAP2-I -9.0<br>PfAP2-O5 -11.1 | 0.16 μM | 93% |
| TCMDC-139888 | | PfAP2-I -9.2<br>PvAP2-I -8.2<br>PfAP2-O5 -8.9 | 1.13 μM | 84% |
| TCMDC-140881 | | PfAP2-I -9.8<br>PvAP2-I -8.2<br>PfAP2-O5 -10.1 | 0.84 μM | 20% |
| TCMDC-124223 |  | PfAP2-I -11.3<br>PvAP2-I -8.9<br>PfAP2-O5 -10.8 | nd | nd |
| TCMDC-125666 |  | PfAP2-I -10.3<br>PvAP2-I -9.3<br>PfAP2-O5 -11.7 | nd | nd |
| TCMDC-142297 |  | PfAP2-I -4.3<br>PvAP2-I -4.2<br>PfAP2-O5 -3.2 | nd | nd |

Our favorite compound is shown in green. Four other lead compounds are shown in black. The two selected compounds exhibiting a tendency to dissociate from the protein during MD simulations are shown in blue. The negative control is indicated in red.
The optimal docking energy score of each compound is given for the ACDC domain of each of the 3 proteins. When available (nd = not determined), the inhibition characteristics of these selected TCAMS compounds are indicated (Gamo *et al.*, 2010).

**Figure 5:**
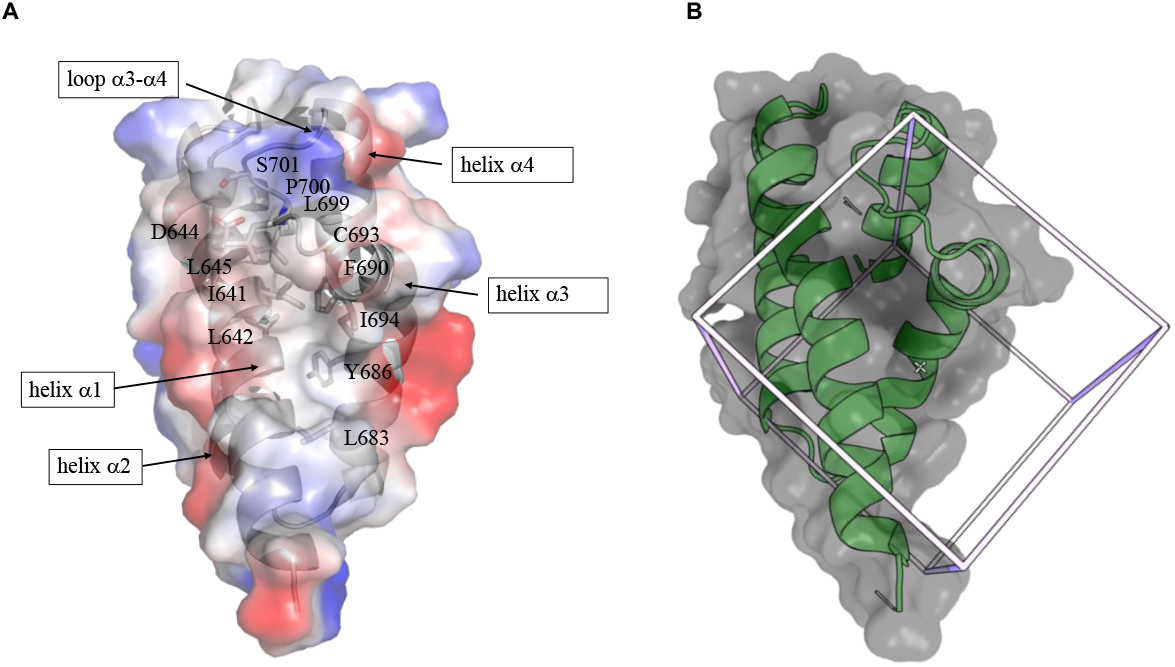
Targeted hydrophobic pocket. **A -** Electrostatic surface of the ACDC domain of PfAP2-O5 shown with positively and negatively charged residues colored in blue and red, respectively. Hydrophobic regions are white. Conserved residues are highlighted as sticks and labeled. **B –** Screening box used for docking highlighted on the structure of the PfAP2-O5 ACDC domain shown in the same orientation as in panel A.

The 7 selected ligands and a negative control (**Figure 6A** and **Table 2**) were submitted to molecular dynamics (MD) simulations for 100 ns (**Sup. Videos 1, 2 and 3**) (**Sup. Figure 5**). Stability throughout the MD simulations was monitored by tracking the minimum distance between the ligands and the pocket, thus assessing the ability of the ligand to remain stably associated with the protein surface and, consequently, their potential as ACDC binders. We further evaluated each ligand thorough visual analysis of their dynamic behavior. As expected, the negative control, TCMDC-142297, disengaged from the pocket early in the simulations and remained detached across all three systems in most of the trajectories. TCMDC-124223 also exhibited a tendency to dissociate and not rebind, while the related molecule TCMDC-125666 only maintained its docked position with PfAP2-O5. However, five of the selected molecules, TCMDC-124284, TCMDC-134543, TCMDC-137354, TCMDC-139888 and TCMDC-140881 exhibited commendable stability throughout the MD simulations, signifying stable interaction with the three ACDC domains. The standout performer was TCMDC-134543, which bound robustly to all three domains (**Figure 6B**), underscoring its potential as a versatile ligand of several ACDC domains.

**Figure 6:**
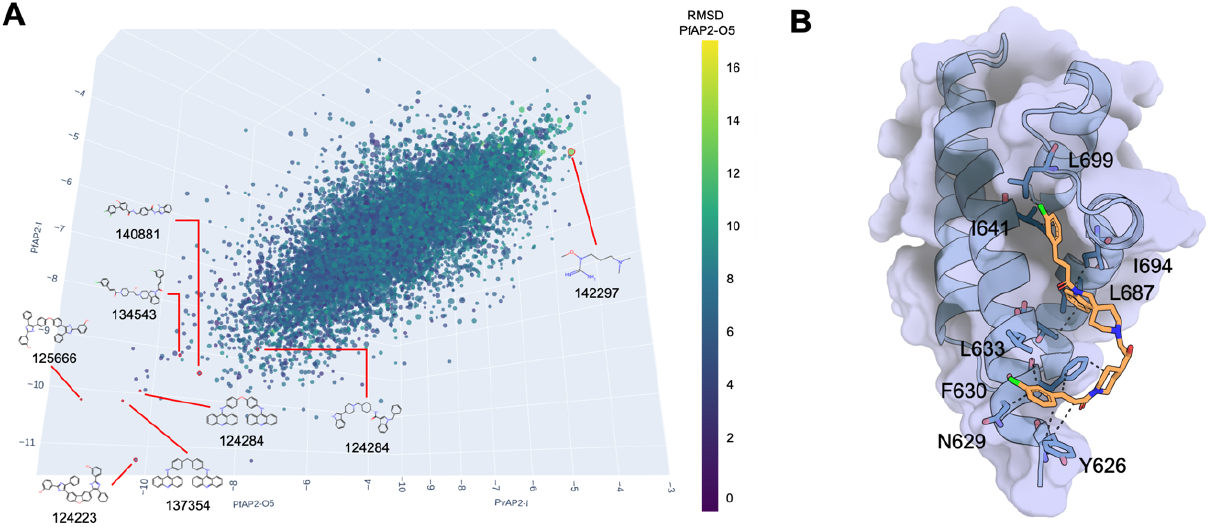
Docking results for the ACDC domains of PfAP2-I, PvAP2-I and PfAP2-O5. **A** - 3D scatter plot showing the docking energy scores (kcal/mol) for the 3 systems with highlight on the 7 selected compounds. TCMDC-142297, which has an unfavorable affinity score but showed minimal distance to the pocket and low average RMSD, was also selected as negative control or MD simulations. The points are colored according to the RMSD (Å) observed in the PfAP2-O5 system from blue (low values) to yellow (high values) whereas the size of the points correlates with RMSD values in the PvAP2-I system. **B** – Selected MD pose for the standout ligand TCMDC-134543 in PvAP2-O5 conserved pocket. Dashed lines represent hydrophobic interactions with residues Tyr626, Asn629, Phe630, Leu633, Thr634 and Ile641 from helix α1 and residues Leu687, Ile694 and Leu699 from helix α3 and loop α3-α4.

## Discussion and Conclusion

Here we show that despite low sequence conservation and position within the proteins (N-terminal versus C-terminal), the ACDC domains from two distinct *P. falciparum* proteins adopt a similar structure. The C-terminal helix swapping observed in the N-terminal ACDC domain of PfAP2-I could be due to the absence of an adjacent folded domain in the crystallized protein fragments and there is currently no evidence that this swapping exists in the full-length protein. Because the PfAP2-I full-length protein contains long disordered poly-asparagine segments, neither experimental structure of the full-length protein nor reliable AlphaFold2 models could be obtained to verify this hypothesis. However, because the ACDC domain of PfAP2-O5 is at the C-terminus and therefore no missing extension can impact the observed α4 position, we consider the PfAP2-O5 conformation as the most probable fold of the ACDC domains. This left-handed orthogonal four-helix bundle is characterized by a kink in helix α3, which constitutes a signature for this new fold. Interestingly, this new fold is predicted to be present in the ACDC domains of all Apicomplexan ApiAP2 proteins despite their low sequence conservation.

Our structural characterization of the ACDC domain does not reveal a function for this domain. Future studies focused on the molecular function of the ACDC domain should shed light on its role in ApiAP2-based transcriptional regulation of gene regulation in Apicomplexan parasites. However, the propensity to make strong contacts in crystal packing suggests that ACDC domains may be involved (i) in interactions between protein partners and the ApiAP2 proteins, or (ii) in dimerization, in line with the tandem occurrence of the TGCA DNA-sequence motif recognized by PfAP2-I (Santos *et al*., 2017). We propose that, within the context of the full-length protein, the ACDC domain could be involved in protein-protein interactions. Interestingly, an ACDC domain has recently been identified in a *Plasmodium* protein (AP2R-2) that lacks functional AP2 domains (Yuda *et al*., 2021). In *P. berghei*, PbAP2R-2 has been shown to interact with the ApiAP2 protein PbAP2-FG2, which also contains an ACDC domain, to form a transcriptional repressor complex (Nishi *et al*., 2023), supporting the hypothesis that the ACDC domain mediates protein-protein interactions.

Given that the ACDC domain adopts a never-before-seen fold and it is exclusively found in Apicomplexa (https://www.ebi.ac.uk/interpro/entry/pfam/PF14733/taxonomy/uniprot/#sunburst), it can be considered as a new therapeutic target. We thus tested the ability of antimalarial inhibitors from the TCAMS drug library to bind ACDC domains. Our *in silico* analysis identified 5 lead compounds, including one, TCMDC-134543, with the potential to target a conserved hydrophobic pocket of several ACDC domains. However, because TCMDC-134543 and TCMDC-124284 are reported, respectively, as inhibitors of human ion channels and ErB2 protein kinase (Gamo *et al*., 2010), and all compounds display HepG2 growth inhibition (**Table 2**), these inhibitors require optimization to improve their specificity and selectivity. Further experiments should also confirm interaction of the ACDC domains with the selected molecules. Nevertheless, our work provides a new therapeutic opportunity for the development of pan-Apicomplexan inhibitors targeting ACDC domain-containing ApiAP2 proteins, which are known to play essential roles in several stages of parasite development.

## Acknowledgements

This work benefited from the crystallization platform of I2BC, supported by the French Infrastructure for Integrated Structural Biology (FRISBI) [ANR-10-INSB-05-05]. We acknowledge SOLEIL for provision of synchrotron radiation facilities, and we would like to thank the staff of beamlines Proxima-1 and Proxima-2A for assistance. JMS team is supported by a CNRS ATIP-Avenir grant. MLB is supported by a doctoral fellowship from the French Ministry of Higher Education and Research *via* the doctoral school Therapeutic Innovation n°569. ML was funded through NIH/NIAID R01AI125565 and PRW was supported by an International Postdoc grant from the Swedish Research Council. This work was also supported by a seeding grant from the MICROBES center for interdisciplinary microbial sciences at the Paris-Saclay University attributed to SN. Computational work was performed using HPC resources from GENCI–IDRIS (Grant AD010714400). TT was granted a postdoctoral fellowship by the ANRS-MIE (ECTZ189696)

