## Supplementary Figures and Tables for "Structural characterization of the ACDC domain from ApiAP2 proteins of the malaria parasite"

### Supplementary data

#### Supplementary figures

**Sup. Figure 1:** Signature sequence of the ACDC domain. The LOGO sequence is calculated based on 1016 aligned sequences of ACDC domains in proteins from various Apicomplexa (Plasmodium, Toxoplasma, Cyclospora, Eimeria…). Occupancy, Insertion probability and insertion length are indicated for each position (https://www.ebi.ac.uk/interpro/entry/pfam/PF14733/logo/).

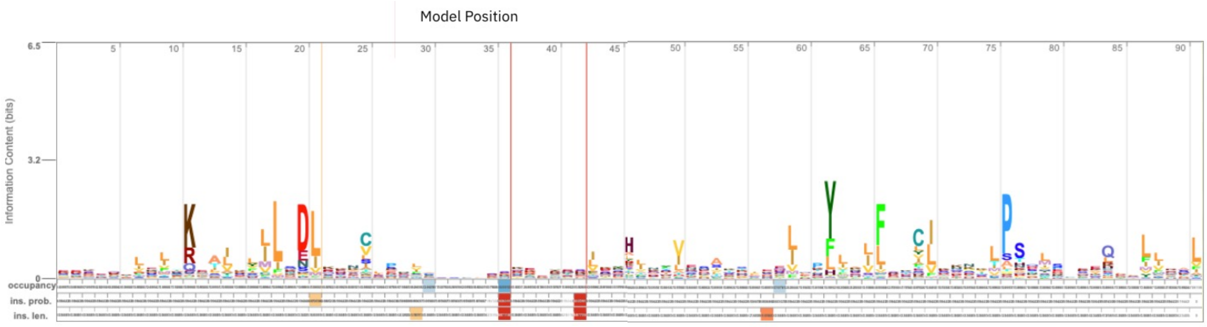

**Sup. Figure 2:** Size-exclusion chromatography (SEC) profile of the PfAP2-O5 ACDC domain during the last step of the purification protocol on a Hiload 16/60 Superdex 75 prep grade column. According to the calibration of the column, the main peak is compatible with a monomeric structure (13 kDa). Arrows indicate elution volumes of protein standards with their corresponding molecular weights.

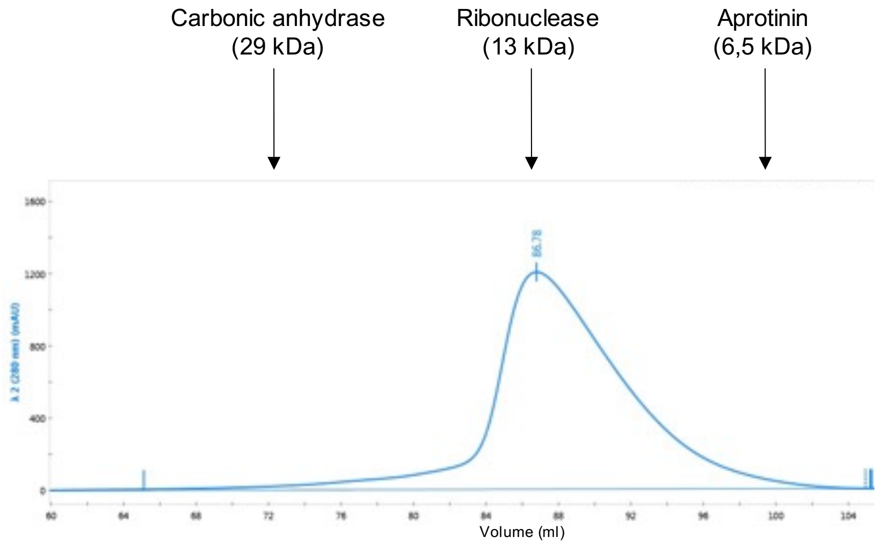

**Sup. Figure 3:** Omit map of loop α3-α4. Electron density map calculated with the final model deleted for loop α3-α4 (residues 120-130). The (2Fo-Fc) map is shown in blue contoured at 2σ. The positive and negative difference maps (Fo-Fc) contoured at 3σ are shown in green and red, respectively. The positive density (in green) indicates features present in the data that are not accounted for by the model whereas the negative density (in red) indicates parts of the model that are not supported by the data. The final structure of the protein is shown as ribbon colored by spectrum.

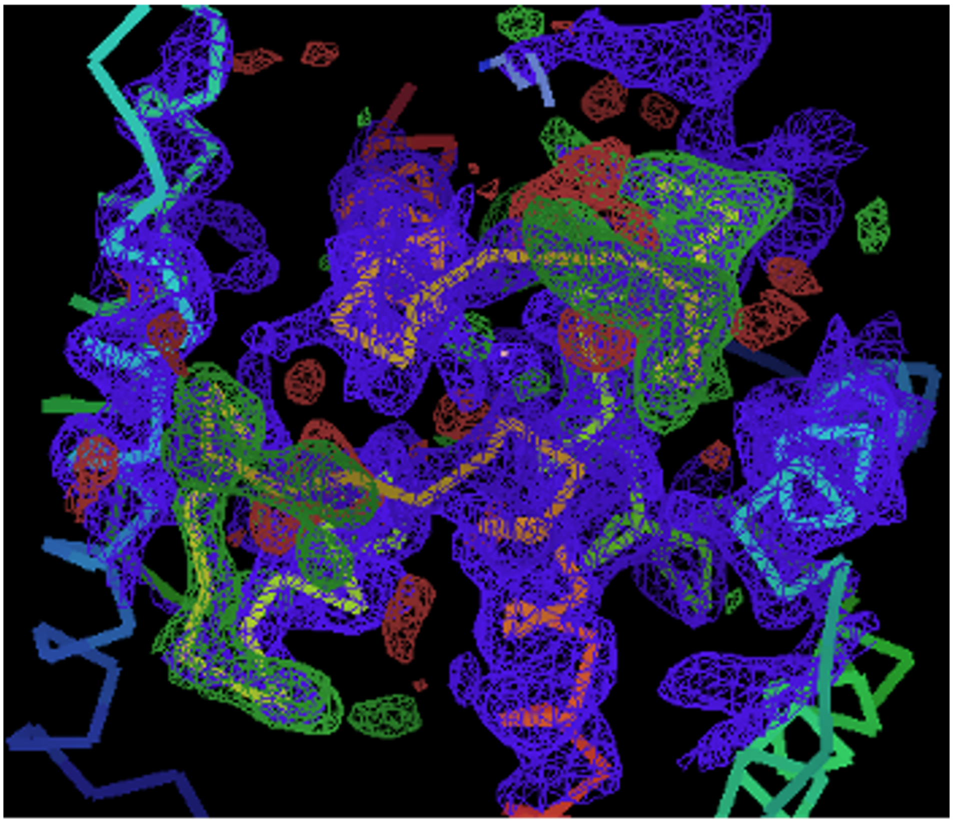

**Sup. Figure 4**: Histogram displaying the distribution of energy scores (in kcal.mol$^{-1}$) of each ligand for all poses in the three systems (PfAP2-I, PvAP2-I and PfAP2-O5). Based on the 40599 datapoints, the distribution is centered around -6.7 kcal.mol$^{-1}$. We kept 7 ligands with the energy score bellow -8.2 kcal.mol$^{-1}$ and one ligand above -4.5 kcal.mol$^{-1}$as negative control.

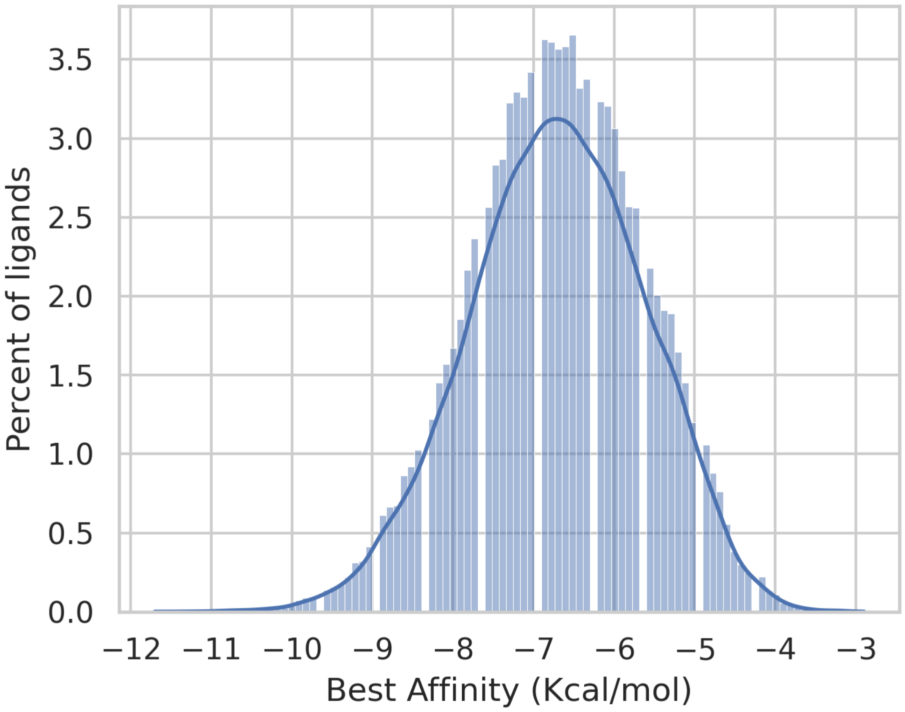


**Sup. Figure 5**: Docking pose for all selected compounds in PfAP2-I, PvAP2-I and PfAP2-O5. The proteins are shown as cartoon colored in green with their surface in transparency. The targeted hydrophobic pocket is highlighted in orange. The docked ligands are shown in sticks colored per atom type (carbon in gray, nitrogen in blue, oxygen in red, fluor in green).

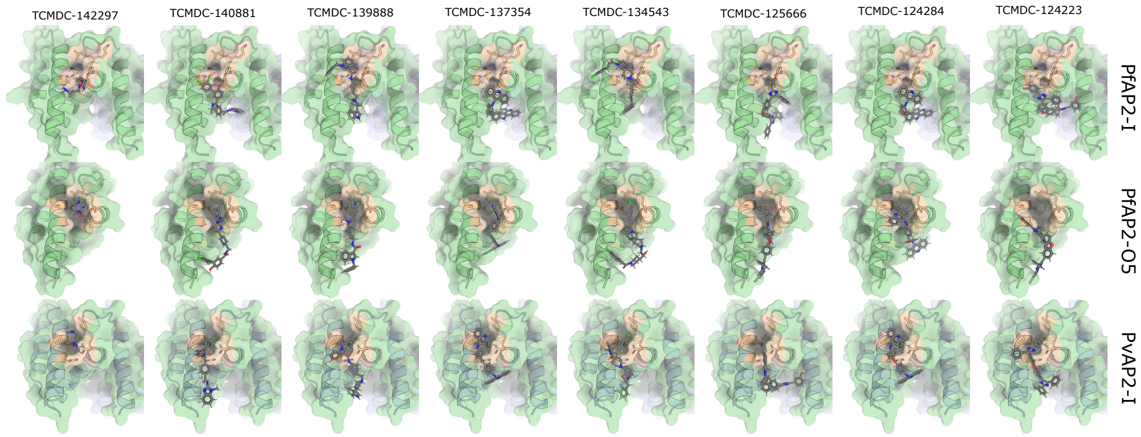

#### Supplementary tables:

**Sup. Table 1**: Primers used for cloning the ACDC domains of PfAP2-O5, PfAP2-I and PvAP2-I.

| **Name** | **Sequence** |
|---|---|
| Fw_PfACDC_AP2-O5 | AACTTCATCGACCTGACCAG |
| Rv_PfACDC_AP2-O5 | CTCGAGCAGCTTGTCCAGGTAGAAC |
| Fw_PfACDC_AP2-I | GCCGCCGCCCATGGATGGAGACATTGGTAAAC |
| Rv_PfACDC_AP2-I | GCCGCCCTCGAGTTATTTTTCAAACTGCGGATGGCTCCATGCGTTCTGTTGAGAATC |
| Fw_PvACDC_AP2-I | TTTTTCCATGGGAAGCCTGAACGAACAC |
| Rv_PvACDC_AP2-I | TTTTTCTCGAGTCATTTTTCAAACTGCGGATGGCTCCAAGCGTTGATGCTGAT |

**Sup. Table 2 :** Parameters of the screening boxes used for molecular docking

| | **PfAP2-I** | **PvAP2-I** | **PfAP2-O5** |
|---|---|---|---|
| Size (x,y,z) | 23,22,20 | 20,29,20 | 23,22,20 |
| Spacing | 1 | 1 | 1 |
| Center grid (x,y,z) | 23.752, 24.939, -2.727 | 23.752, 24.939, -2.727 | 23.752, 24.939, -2.727 |
